# DMD Human iPSC-derived skeletal muscle cells recapitulate selective calcium dysregulation pathways

**DOI:** 10.1101/2024.01.20.576393

**Authors:** Arnaud Delafenêtre, Charles-Albert Chapotte-Baldacci, Léa Dorémus, Emmanuelle Massouridès, Marianne Bernard, Matthieu Régnacq, Jérôme Piquereau, Aurélien Chatelier, Christian Cognard, Christian Pinset, Stéphane Sebille

**Affiliations:** PRETI laboratory, University of Poitiers, France; CECS, I-STEM, AFM, Corbeil-Essonnes, France

## Abstract

This study investigates the functional characteristics of induced pluripotent stem cell-derived muscle cells (hiPSC-skMCs) from Duchenne muscular dystrophy (DMD) patients, focusing on their regulation of intracellular calcium concentration. DMD, a progressive muscle degenerative disease, arises from mutations in the dystrophin gene and is characterized by elevated intracellular calcium levels, exacerbating disease progression. This work highlights that DMD hiPSC-skMCs demonstrate unique calcium signatures with increased intracellular calcium compared to healthy counterparts. These cells also exhibit both heightened calcium response when stimulated by electrical fields or acetylcholine and more pronounced constitutive calcium entries. While RNAseq data from these cells reaffirmed known dysregulation mechanisms seen in other dystrophin-deficient models, certain pathways like purinergic or store-operated calcium entries did not show disruption in this DMD model. This discrepancy suggests that not all mechanisms observed in animal models may be equally relevant in human cases, pointing towards specific molecular targets that could be more effective for DMD treatment strategies.

## Introduction

Duchenne muscular dystrophy (DMD) caused by mutations in the dystrophin gene, is an X-linked muscle degenerative disease affecting 1.7-2.1 of 10,000 male births (Hoffman et al., 1987; Parsons et al., 2002). In DMD patients, the loss of dystrophin destabilizes the dystrophin glycoprotein complex (DGC), leading to sarcolemma instability and functional deficits (Millay et al., 2009). Dystrophin also anchors dystrophin-associated proteins to the sarcolemma, including neuronal nitric oxide synthase (nNOS) (Brenman et al., 1996). The absence of functional dystrophin results in the degeneration and death of skeletal and cardiac muscles, which are subsequently replaced by fibrous tissue (Desguerre et al., 2009). Currently, there is no curative treatment for DMD.

A significant secondary consequence in DMD is the abnormal elevation of intracellular calcium concentration, (Ca^2+^)_i_, in dystrophin-deficient muscles, which exacerbates disease progression. Calcium is a pivotal biochemical agent in skeletal muscle cell contraction (Ebashi, 1972) and serves as a secondary messenger in various cellular processes across different cell types (Berridge, 1997). Maintaining Ca^2+^ concentration in skeletal muscles is crucial for continuous muscle fiber contractility (Tu et al., 2016). Thus, Ca^2+^ ion movements in skeletal muscles involve intricate interactions among various regulated subsystems (Bravo-Sagua et al., 2020). The intracellular calcium overload seen in DMD is believed to influence its pathogenesis (Uchimura et al., 2021), suggesting that modulating Ca^2+^ handling mechanisms might offer a promising therapeutic approach for DMD. Studies have shown that the sarcolemma, sarcoplasmic reticulum, and mitochondria, through diverse molecular pathways alterations, contribute to the persistent elevation of (Ca^2+^)_i_ levels (for review, see Mareedu et al. 2021). Elevated Ca^2+^ levels activate Ca^2+^-dependent proteases and phospholipases, causing muscle necrosis, impairing mitochondrial function, and increasing reactive oxygen species (ROS) production. These collective alterations result in muscle atrophy and contractile dysfunction.

Our current understanding of DMD pathophysiology primarily stems from studies using mouse models (mdx and its variants) that lack dystrophin and often exhibit a milder DMD phenotype (McGreevy et al., 2015). However, these models have led to only a few therapeutic advances, highlighting the need for human-centric research models. Notably, studies investigating the role of calcium in muscle cell differentiation using human induced pluripotent stem cells (hiPSCs) are relatively scarce. Utilizing hiPSCs in research offers a unique platform to explore muscle cell differentiation mechanisms within a human context, potentially enhancing our comprehension of muscle disorders in humans. Therefore, establishing a disease model that accurately reflects clinical symptoms or molecular pathogenesis is crucial.

Previously used Duchenne muscular dystrophy (DMD) hiPSC lines from patients with dystrophin gene mutations (Mournetas et al., 2021) showed two main outcomes in transcriptome and miRnome comparisons with healthy cells. First, hiPSC differentiation accurately mirrors the developmental path from mesoderm to skeletal muscle. Second, this process yields myotubes displaying DMD phenotypes (Mournetas et al., 2021). Based on these insights, the current study hypothesized that muscle cells derived from DMD patient hiPSCs may replicate the calcium dysregulation seen in DMD. Consequently, we explored the functional behaviors of both healthy and DMD hiPSC-derived muscle cells, particularly their management of intracellular calcium levels under both resting conditions and when subjected to electrical and pharmacological stimuli.

## Results

### Morphology of human induced pluripotent stem cell-derived muscle cells (hiPSC-skMCs)

hiPSCs from DMD patients and healthy individuals were generated and grown as described previously (Massouridès et al., 2015; Mournetas et al., 2021). These cells were subjected to a standardized differentiation protocol. Thawed at myoblast stage, cells proliferated for 3 days, which corresponds to the myoblast stage (D3). Myoblasts were incubated for 4 days in a medium inducing cell differentiation (SKM03) into elongated and plurinucleated myotubes (D7). Figure 1 describes morphology of muscle cells derived from healthy hiPSC-skMCs during amplification of myoblasts (D3) (A) and during fusion process in myotubes (D7) (B). In D7, cells were aligned and displayed elongated shapes. Fluorescent staining of nuclei and α-actinin (Figure 1C) confirmed both the fusion in displaying several nuclei per cell and the formation of striation patterns in elongated cells. Myosin heavy chains labelling with MF-20 antibody demonstrated their presence in striated myotubes (Figure 1D). DMD hiPSC-skMCs myotubes displayed also α-actinin striations at the myotube stage (Figure 1E) and electron microscopy images of such cells allowed to detect the establishment of the alignment of myosin filaments in myotubes, without striations (Figure 1F). Both DMD and healthy myotubes could be sustained in culture for extended periods. However, beyond day 7, the cells exhibited pronounced contractile activities, causing them to detach from the substrate. This detachment hindered our ability to conduct most stimulation experiments.

**Figure 1:**
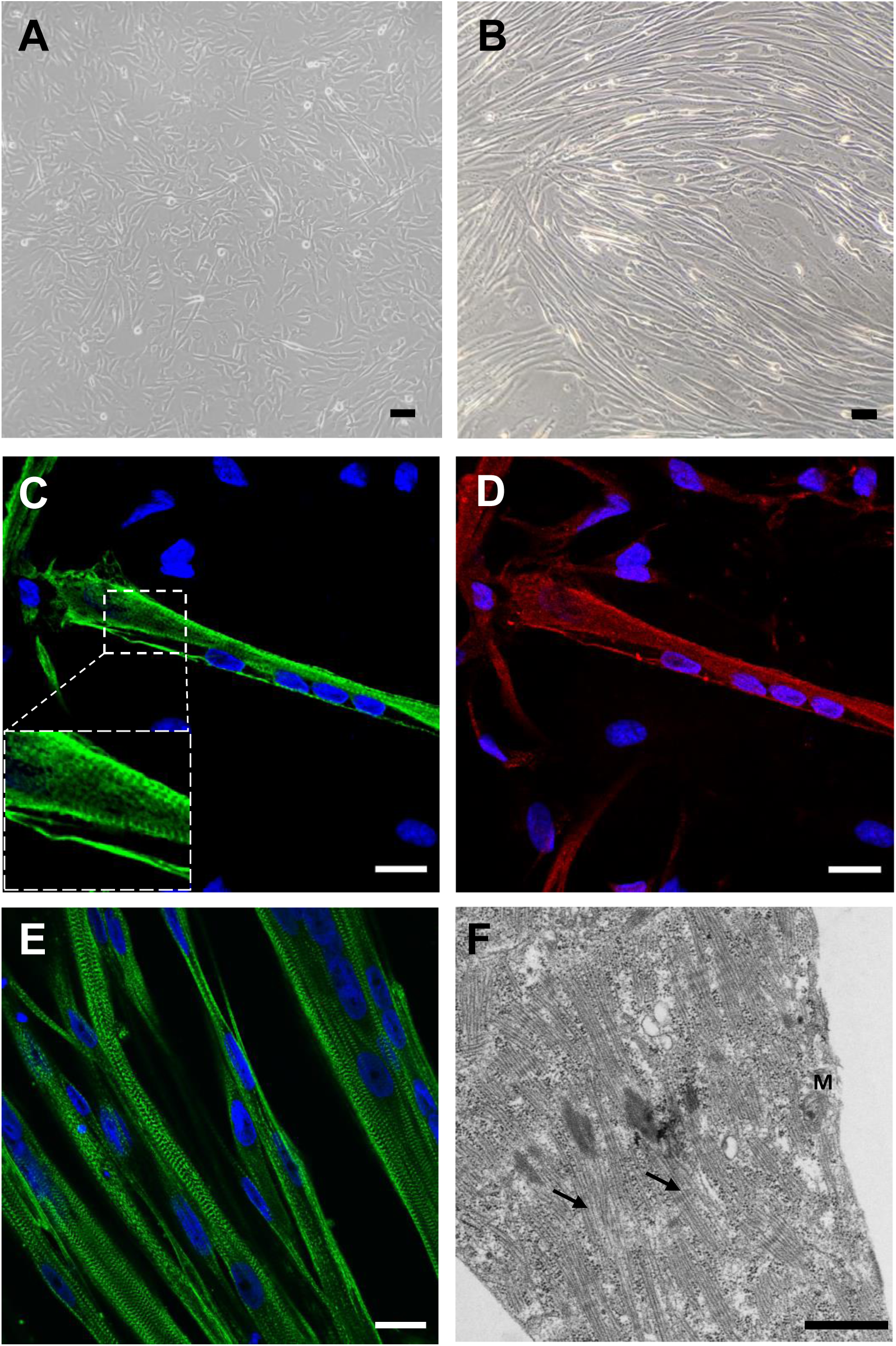
Morphology and ultrastructure of human induced pluripotent stem cell-derived muscle cells (hiPSC-skMCs). **A and B:** Transmission images of hiPSC-derived healthy (M180) myoblasts (D3) (**A**) and after differentiation (myotubes: D7) (**B**). Scale bar: 50 µm. **C and D**: Immunostaining of healthy hiPSC-derived myotubes (D7) with α-actinin in green (**C**) and myosin heavy chain in red (**D**). Nuclei are labeled in blue (Dapi); scale bar: 20 μm. **E:** Immunostaining of DMD (M418) hiPSC-derived myotubes with α-actinin in green; scale bar: 20 μm. **F:** Electron microscopy images of hiPSC-derived myotubes (M398) (arrows indicate myosin filaments). Scale bar: 2 μm.

### Resting calcium levels and membrane potential values in healthy and DMD hiPSC-skMCs

Intracellular calcium concentrations ([Ca^2+^]_i_) have been determined, at rest, in both healthy and DMD hiPSC-skMCs lines and at both D3 and D7 stages, (Figure 2A). Intracellular calcium concentrations were found to be close to normal values (Gehlert et al., 2015) in healthy cells. For DMD cells, [Ca^2+^]_i_ were found elevated at D3 stage. At D7 stage, no significant differences were found between healthy and DMD cells. Resting membrane potential values have also been measured with the microelectrode method in our hiPSC-skMCs cell lines at three different maturation stages (D3, D7 and D10: Figure 2B and S1). Resting membrane potential of a differentiating muscle cell allows to evaluate its excitability properties and therefore its ability to activate excitation-contraction coupling mechanism. In D3 stage, cells displayed membrane potential values of - 9.0 ± 1.0 mV for healthy cells and −8.1 ± 0.6 mV for DMD cells, meaning that, at this stage, cell membranes are weakly polarized (Figure 2B), as often seen in proliferating cells. Polarization of membranes increased for both cell lines in D7(−24.0 ± 1.9 mV for healthy cells and −19.2 ± 3.4 mV for DMD cells) and in D10 conditions (−32.8 ± 3.0 mV for healthy cells and −30.5 ± 3.7 mV for DMD cells) in which most cells formed myotubes. These findings are consistent with previous observations regarding muscle cell polarization during the maturation process (Fennelly and Soker, 2019). Here, we can assume that hiPSC-derived muscle cells were subsequently able to generate the physiological machinery, comprising various channels and pump proteins, which facilitate ion movement across the lipid bilayer and establish negative resting potential values. No significant difference in resting membrane potential values was found between healthy and DMD cells.

**Figure 2:**
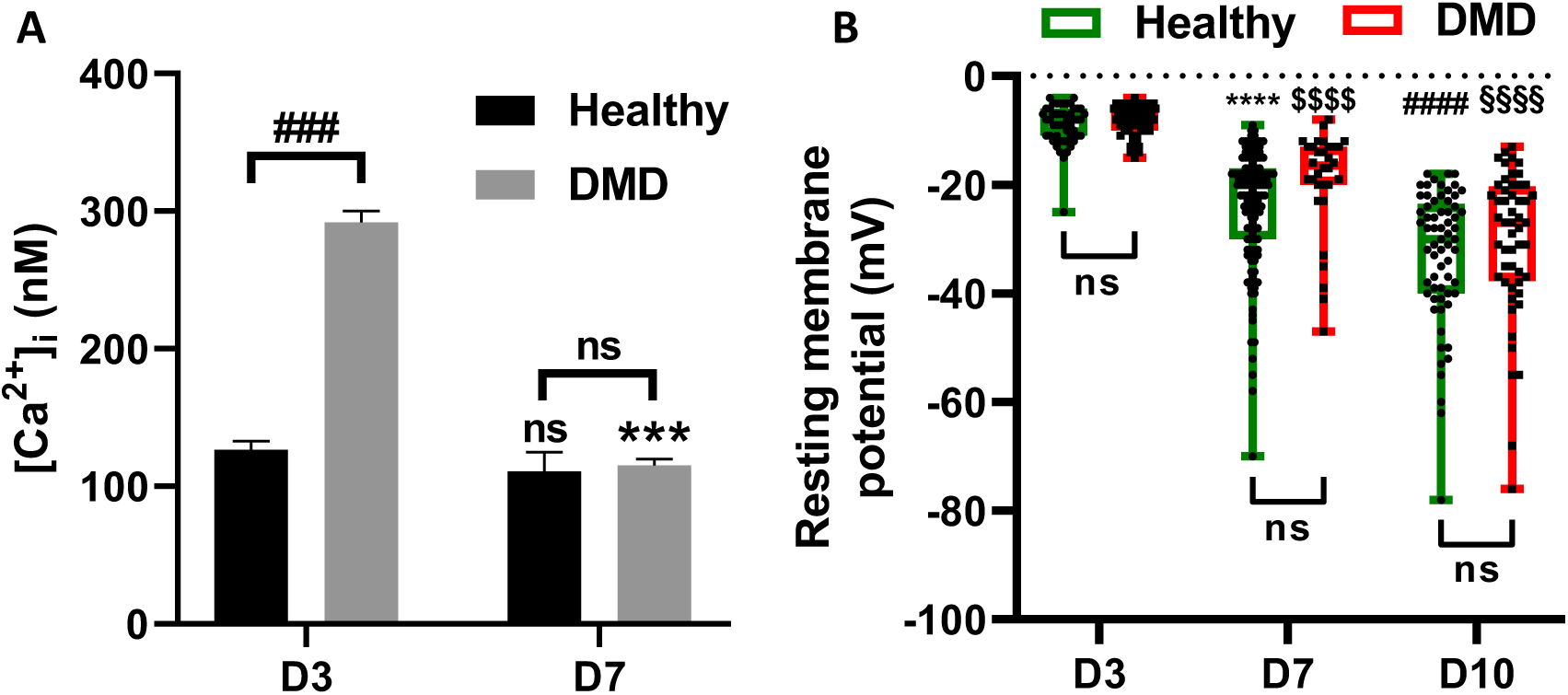
Resting calcium levels and membrane potential values in healthy and DMD hiPSC-skMCs. **A:** Intracellular calcium concentrations determined using fura-2 dye and ratiometric method in healthy (M180) and DMD (M202) hiPSC-skMCs lines at D3 and D7. (Results shown as mean ± SEM; Healthy: D3, n=56; D7, n=80. DMD: D3, n=97; D7, n=92). * indicates the significant difference between D3 and D7 cells, and # between healthy and DMD. *** and ###: *p* < 0.005, Kruskal Wallis test with Dunn’s multiple comparisons test). **B:** Resting membrane potential values measured with the microelectrode method in both healthy (M180 and M194) (green) and DMD (M202 and M418) (red) hiPSC-skMCs at three different maturation stages, D3, D7 and D10. (* indicates the significant difference between D3 and D7 healthy cells. # Indicates the significant difference between D7 and D10 healthy cells. $ indicates the significant difference between D3 and D7 DMD cells. § Indicates the significant difference between D7 and D10 DMD cells Healthy: D3, n= 50; D7, n=133; D7+3, n=60. DMD: D3, n=66; D7, n=32; D7+3, n=52. **** and ####: *p* < 0.001, Kruskal Wallis test with Dunn’s multiple comparisons test.)

### Spontaneous calcium increases in differentiating hiPSC-skMCs

During in vitro maturation, skeletal muscle cells can exhibit spontaneous calcium activities, i.e. local or global calcium increase events (Balghi et al. 2006b; Chapotte-Baldacci et al. 2022). To record large and global activities, currently termed calcium oscillations, calcium imaging has been performed with confocal microscopy in DMD and healthy hiPSC-skMCs cell lines (Figure 3A-B). Recording of spontaneous global calcium oscillations revealed multiple event profiles with varying durations and frequencies (Figure 3B). Each of these events has been analyzed to calculate the amplitude and three kinetic parameters (Time to Peak: TTP, Duration and Recovery Time: RT) allowing to compare spontaneous events in healthy and DMD cells in the two differentiation stage myoblasts (D3) and myotubes (D7). Results showed that the amplitude of calcium events increased with cell differentiation in both cell types (Figure 3C). Moreover, DMD cells at D7 showed a higher significant amplitude (2.99 ± 0.06) than healthy cells (2.29 ± 0.08). Analysis of kinetic parameters displayed a decrease of events duration, associated with a reduced TTP, in DMD cells at D7 cells as compared to events from healthy cells (Figure 3 D-E), with no changes in RT (Figure 3F).

**Figure 3:**
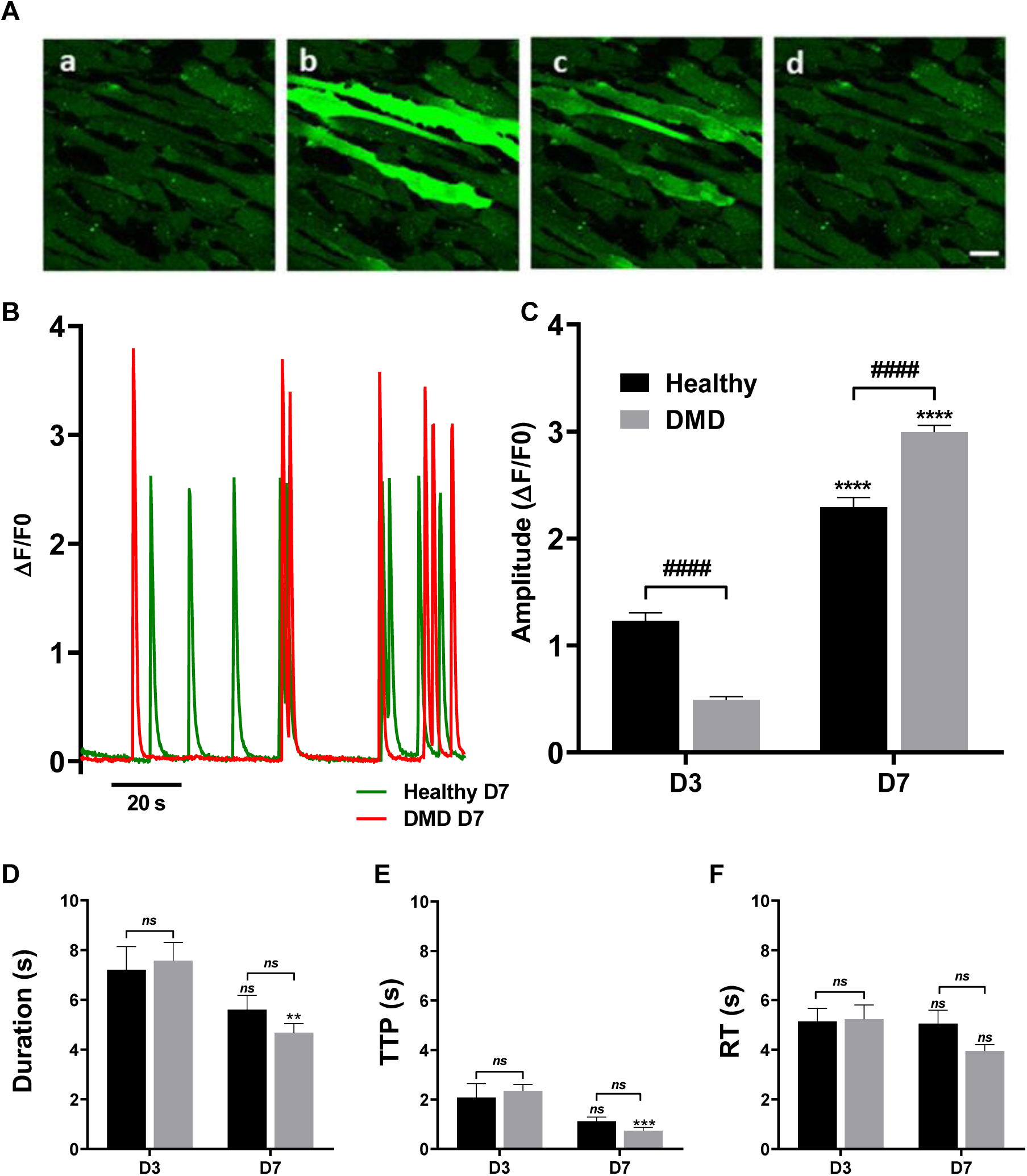
Spontaneous calcium oscillations in healthy and DMD hiPSC-skMCs. **A:** Image series (a, b, c, d) of fluo-8 fluorescence during spontaneous calcium oscillations in healthy (M180) hiPSCs (5 fps). **B:** Examples of calcium kinetics in hiPSCs-derived myotubes in healthy (M180) and DMD (M202) cells at D7. Amplitude **(C)**, Duration **(D)**, Time to peak (TTP: **E**) and Recovery time (RT: **F**) of calcium events in healthy and DMD hiPSCs-derived cells at D3 and D7. (Healthy: D3, n=102; D7, n=275. DMD: D3, n=168; D7, n=207. * Indicates the significant difference between D3 and D7. # indicates the significant difference between healthy and DMD lines. **: *p* < 0.01; ***: *p* < 0.005; **** and ####: *p* < 0.001, Kruskal Wallis test with Dunn’s multiple comparisons test.)

hiPSC-skMCs also displayed a large amount of highly localized calcium increases (Figure 4) that were demonstrated to be mainly due to local calcium releases from sarcoplasmic reticulum in developing muscle cells (Balghi et al., 2006). These local events were recorded and analyzed through two different methods. First, release site density was calculated through the computation of the standard deviation on image series of fluo-8 loaded cells (Figure 4A-B). Even if the number of release sites seemed not to depend on the differentiation stage in healthy cells, it was found significantly increased in DMD D7 cells (Figure 4B). The morphology of these events has then been characterized and fluo-8 images were recorded in a line scan mode (x, t) taking fluorescence variations along a line as a function of time (Figure 4C). Local events were found longer along the time in DMD D7 cells as compared to healthy cells, regarding their elevated durations and RT (Figure 4E and G respectively). All these data demonstrated that spontaneous calcium activity revealed an increase in the amount of calcium released during calcium oscillation throughout differentiation.

**Figure 4:**
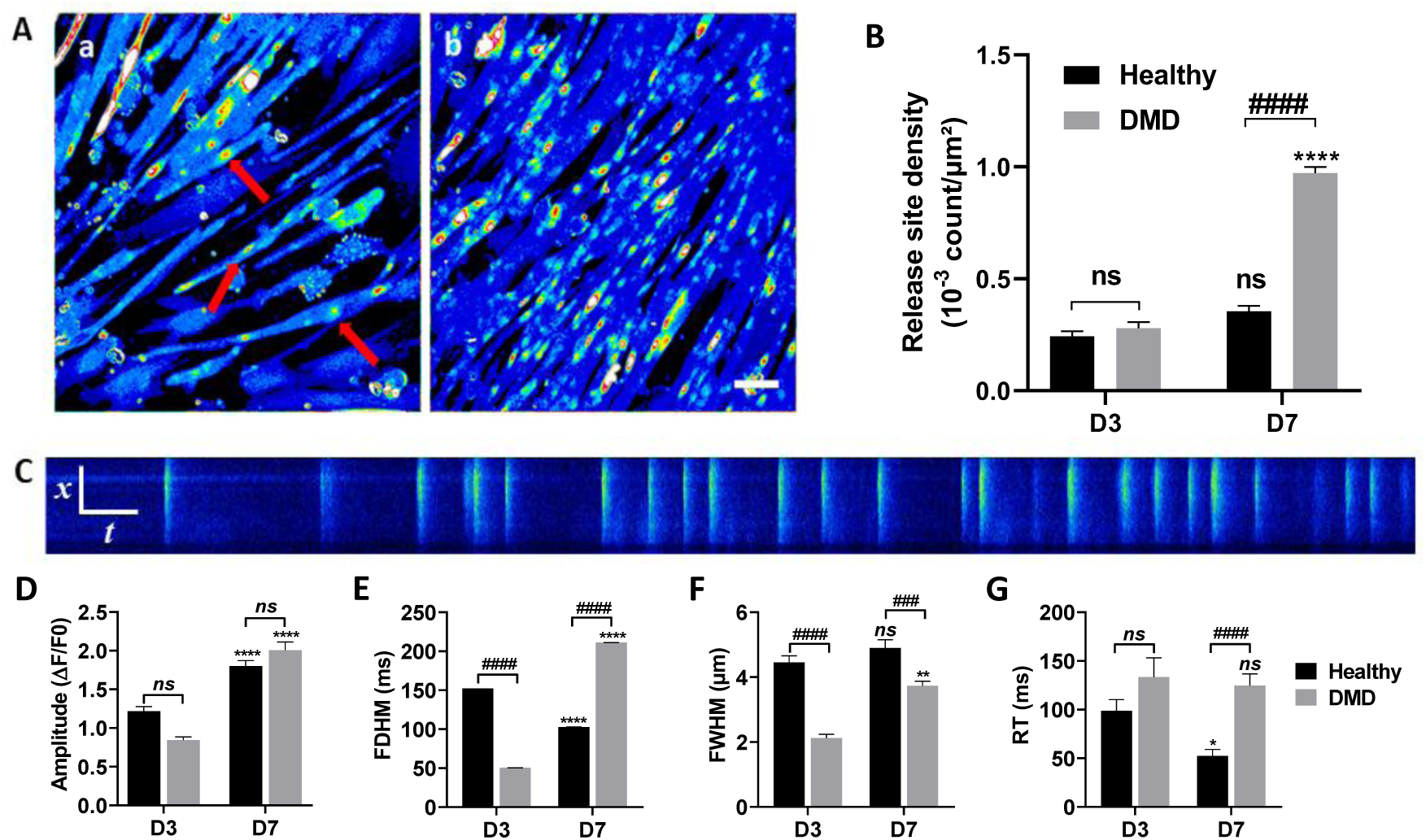
Spontaneous local releases of calcium in healthy and DMD hiPSC-skMCs. **A:** Standard deviation images of calcium fluorescence recording (1 min) showing sites of calcium release (red arrows) identified in a culture of healthy (M180) (a) and DMD (M202) (b) hiPSC-CMs at D7. Scale bar: 20 μm. **B:** Release site density in healthy and DMD hiPSCs-derived myoblasts and myotubes (data are shown as mean ± SEM. Healthy: D3, n=19; D7, n=20. DMD: D3, n=39; D7, n=35. * indicates the significant difference between D3 and D7. # indicates the significant difference between healthy and DMD lines. **** and ####: *p* < 0.001, Kruskal Wallis test with Dunn’s multiple comparisons test). **C:** 2-D images (800*800 pixels) acquired in a line scan mode (x, t images: 1 line/2 ms, 100 pixels) showing fluorescence variations along a line as a function of time. Amplitude (**D**) and kinetic parameters, FDHM (**E**), FWHM (**F**), and RT (**G**) (data are shown as mean ± SEM. Healthy: D3, n=121; D7, n=264. DMD: D3, n=47; D7, n=198. * indicates the significant difference between D3 and D7, and # between healthy and DMD. *: *p* < 0.01, **: *p* < 0.005, **** and ####: *p* < 0.001, Kruskal Wallis test with Dunn’s multiple comparison test).

### Calcium responses of hiPSC-skMCs under electrical stimulation

As it has been demonstrated that hiPSC-skMCs exhibited negative values of membrane potentials, electric field stimulation (EFS) was performed above cell cultures loaded with fluo-8. Calcium increase events were recorded during EFS through calcium imaging with confocal microscopy (Figure 5A-B and video S1). As we anticipated based on the values of membrane potential, not all cells exhibited a calcium response to this stimulation; only 20% of the myoblasts and 35% of the myotubes, regardless of cell line, displayed significant calcium elevation events. Each of these events has been analyzed to calculate amplitude (Figure 5C), area (Figure 5D) and three kinetic parameters (Time to Peak: TTP, Duration and Recovery Time: RT; Figure 5E-G). Results showed that the amplitude of calcium signal increased with cell differentiation in both cell types (Figure 5C). Moreover, DMD cells at D7 showed a higher signal than healthy cells (Figure 5C-D) (1.42 ± 0.07 for healthy cells and 2.13 ± 0.07 for DMD cells). Analysis of kinetic parameters showed significant differences between healthy and DMD lines (Figure 5E-G). Indeed, the duration of calcium release, the time to peak and the rise time of myotubes (D7) were significantly higher in DMD cells as compared to healthy cells (Figure 5E-G). Nifedipine at 1 µM completely blocked calcium increase during electrical field stimulation (Figure S2). These findings imply that calcium increase observed during electrical stimulation are mediated via DHPR channels.

**Figure 5:**
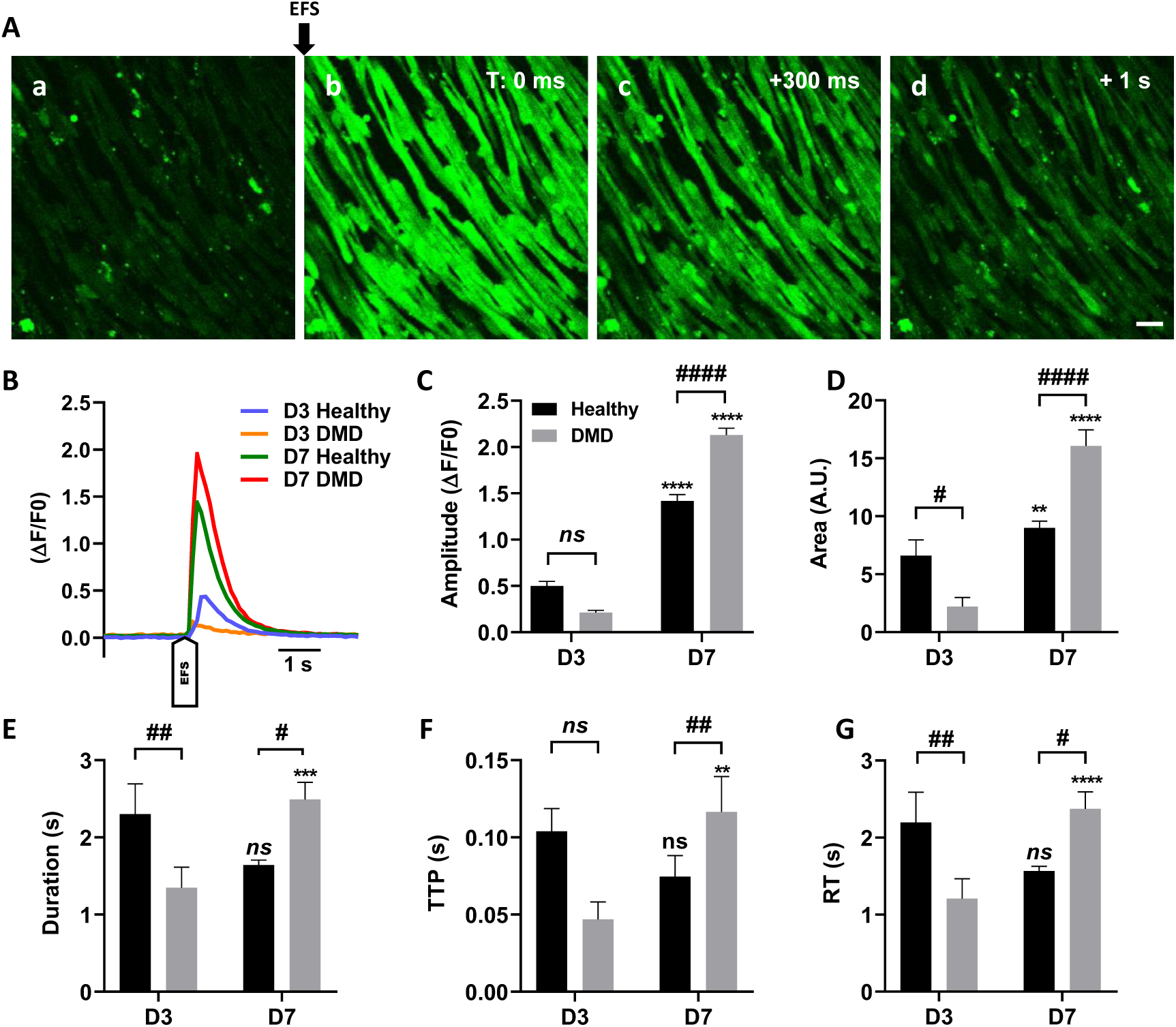
Characterization of calcium responses induced by electrical field stimulation (EFS) in healthy and DMD hiPSC-skMCs. **A.** Representative example of fluo-8 fluorescence measurements before (a) and after (b) an EFS-induced calcium increase in healthy (M180) hiPSC-CMs. Scale bar: 20 μm. **B:** examples of kinetics obtained in each condition. Amplitude (ΔF/F0) (**C**), Area (A.U.) (**D**) as well as kinetic parameters: Duration (**E**), Time to peak (TTP) (**F**) and RT (**G**) of electrically induced calcium increases were determined in healthy (M180) and DMD (M202) hiPSC-CMs (data are shown as mean ± SEM. Healthy: D3, n=60; D7, n=82. DMD: D3, n=35; D7, n=133. * indicates the significant difference between D3 and D7 cells, and # between healthy and DMD. **** and ####: *p* < 0.001, Kruskal Wallis test with Dunn’s multiple comparisons test).

### Pharmacological stimulations

We were then interested to evaluate if our hiPSC-skMCs were able to respond to the acetylcholine, the endogenous neurotransmitter released at the neuromuscular junction, capable of inducing skeletal muscle contraction. Acetylcholine at 100 µM was perfused on hiPSC-skMCs cultures loaded with fluo8-AM and calcium responses were determined through calcium imaging (Figure 6 A). In this case, 52 % of the myoblasts and 97 % of the myotubes, regardless of cell line, displayed significant calcium elevation events. Each of these events has been analyzed to calculate the same parameters as Figure 5. Results show that the amplitude of calcium signal increased with cell differentiation in both cell types (Figures 6B-C and Figure S3). Moreover, DMD cells at D7 displayed a signal with higher amplitude (2.35 ± 0.12) as compared to healthy cells (1.90 ± 0.09). Acetylcholine perfusion was also performed using either an extracellular calcium free solution or nifedipine incubation (10 µM). These two conditions induced a significant reduction of 78 % and 73 %, respectively, in the proportion of responding cells (Figure 6D), demonstrating the involvement of both extracellular calcium and excitation-contraction coupling elements in this acetylcholine-dependent calcium increase. Furthermore, for the few remaining responding cells, calcium signal amplitudes were found significantly reduced in these two inhibitory conditions. (Figure 6E-F).

**Figure 6:**
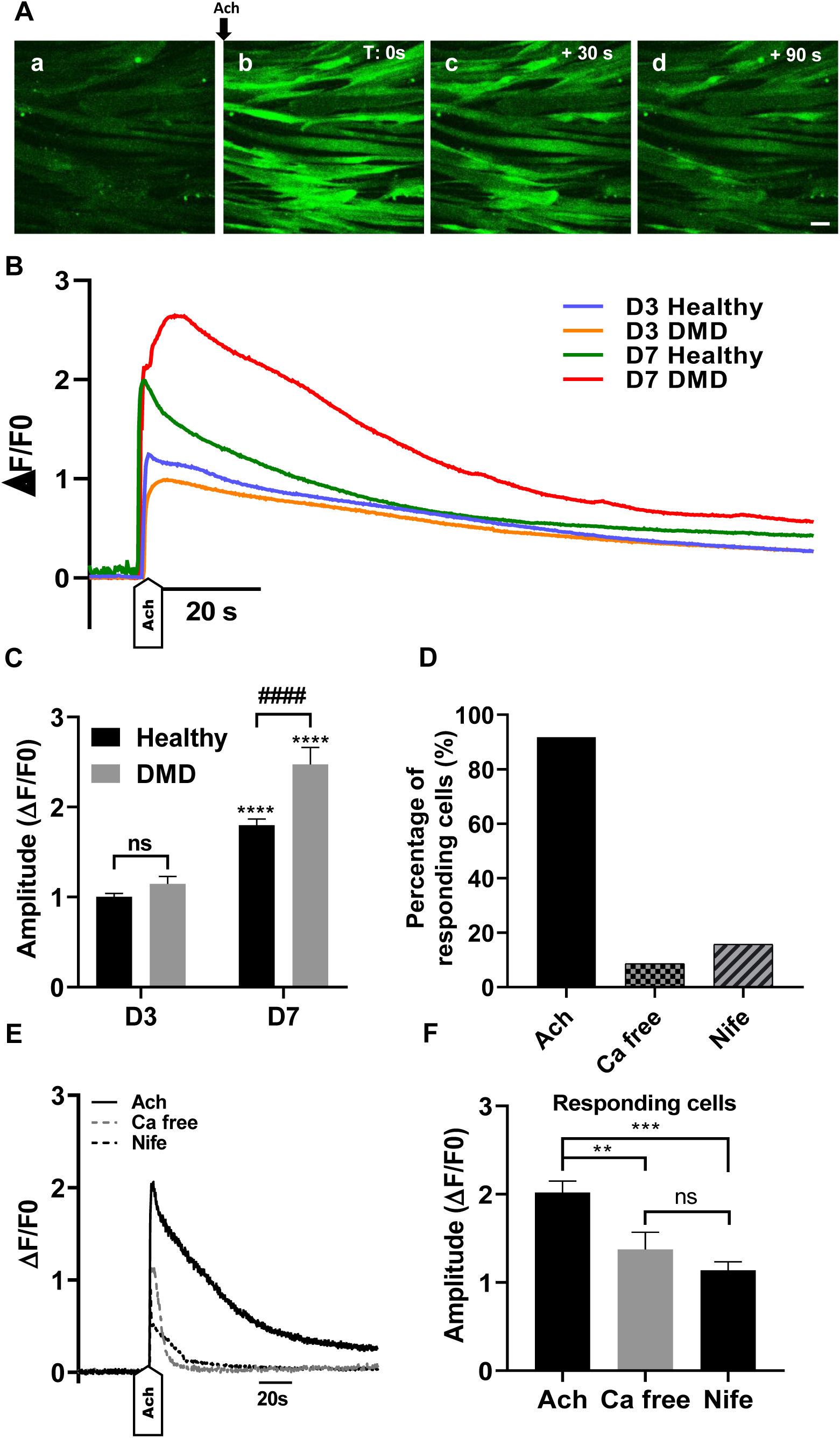
Characterization of calcium responses induced by acetylcholine stimulation in healthy and DMD hiPSC-skMCs. **A:** Example of fluo-8 fluorescence images recorded during acetylcholine perfusion (100 µM) in healthy (M180) and DMD (M202) hiPSC-CMs. Scale bar: 20 μm **B:** Representative kinetics obtained in each condition. **C:** Analysis of the Amplitude (ΔF/F0) of acetylcholine induced calcium increases in DMD and Healthy hiPSCs. **D:** Percentage of responding cells during acetylcholine perfusion in free extracellular calcium and with nifedipine (10 µM). **E:** Kinetics obtained with Healthy hiPSCs myotubes with the amplitude (ΔF/F0) measured for each treatment (**F**). data are shown as mean ± SEM. Healthy: D3, n=16; D7, n=16. DMD: D3, n=16; D7, n=16. * indicates the significant difference between D3 and D7. # indicates the significant difference between healthy and DMD lines. **: *p* < 0.01, ***: *p* < 0.005, **** and ####: *p* < 0.001, Kruskal Wallis test with Dunn’s multiple comparisons test.)

Because elevated activity of P2X7 receptors has been identified in myoblasts from the mdx mouse model of DMD (Rog et al., 2023), purinergic receptors have also been interrogated in our hiPSC-skMCs cultures with BzATP perfusion (100 µM and 300 µM) (Figure S4). BzATP perfusion was also performed on SolC1(-) skeletal muscle cells line, a dystrophin-deficient cell line originating from mouse (Balghi et al., 2006). Results show that BzATP perfusion at 100 µM induced a large calcium increase in cultured SolC1(-) myotubes. However, perfusion of BzATP with similar concentration in cultured DMD hiPSC-skMCs at D7 led to low calcium increase, which moreover involves only a small proportion of cells (around 2% of responding cells). No differences were observed with a 300 µM BzATP perfusion.

### Constitutive and store-operated calcium entries in healthy and DMD hiPSC-skMCs

Constitutive calcium entries have been determined in both healthy and DMD hiPSC-skMCs at the same two stages D3 and D7. Such entries have been previously shown to be elevated in the context of dystrophin deficiency, primarily involving TRP channels (Aguettaz et al., 2016; Sabourin et al., 2009). Mn^2+^ quenching experiments were performed, and the rate of Fura-2 fluorescence quenching was determined as an index of calcium entry (Figure 7). In DMD cells at D7, addition of 0.1 mM Mn^2+^ induced a quenching of the Fura-2 fluorescence, characterizing a constitutive entry of calcium at rest, that is not observed in healthy cells (Figure 7A). In healthy myoblasts and myotubes, only slow constitutive calcium entries have been recorded (−1.5 ± 0.6 %/min at D3 and −1.9 ± 0.2 %/min at D7, Figure 7B). Constitutive calcium entries were found increased in DMD cells at both stages (−21.1 ± 4.0 %/min at D3 and −21.7 ± 2.4 %/min at D7). For both cell lines, no significant differences have been calculated between D3 and D7, indicating that maturation of myoblasts into myotubes did not modify levels of constitutive calcium entries.

**Figure 7:**
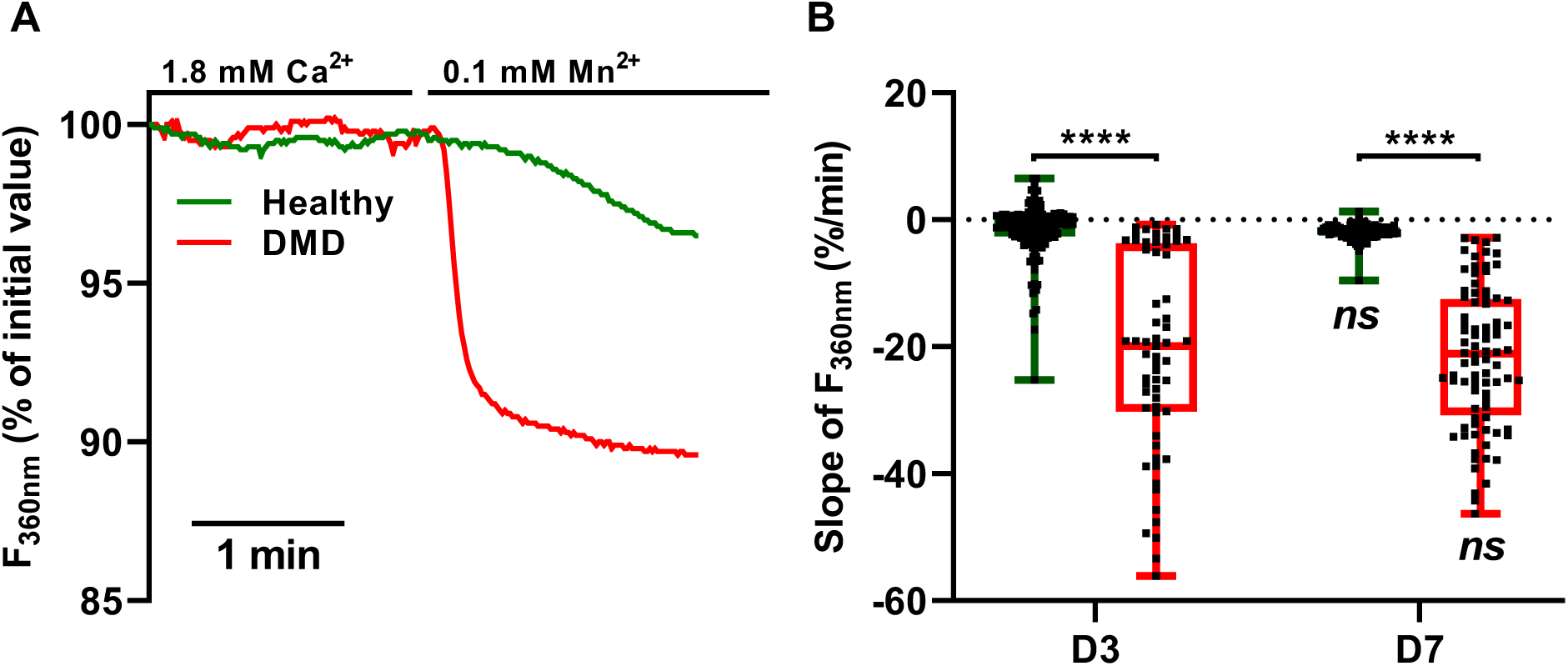
Characterization of constitutive calcium entries (CCE) in healthy and DMD hiPSC-skMCs. **A:** Examples of CCE kinetics in hiPSC-skMCs myotubes in healthy (M180) (green) and DMD (M202) (red) conditions at D7. **B:** CCE measured in both healthy (green) and DMD (red) hiPSC-skMCs cell lines at D3 and D7. (Data are shown as mean ± SEM, healthy: D3, n=161; D7=90. DMD: D3, n=61; D7, n=84. * indicates the significant difference between D3 and D7. # indicates the significant difference between healthy and DMD lines, Kruskal Wallis test with Dunn’s multiple comparisons test).

Store operated calcium entries (SOCEs) have also been determined in both healthy and DMD hiPSC-skMCs at the same two stages D3 and D7. SOCEs have been demonstrated to be involved in calcium dysregulation in several dystrophin deficient models (Goonasekera et al., 2011; Harisseh et al., 2013). Experimental protocols allowing to determine SOCEs have been performed in our models (Figure S5 A), using ionomycin perfusion at the end of each experiment as a positive control for calcium increase slope characterization. Results show that no significant increase of intracellular calcium, originating from SOCs, and characterized with increase slope (Figure S5B) or area (Figure S5C) was observed at D3 in both cell lines. A very low but significant increases were observed at D7 in both cell lines (Figure S5B-C). At this stage, no significant differences were found between healthy and DMD hiPSC-skMCs.

## Discussion

Studying calcium regulation in human induced pluripotent stem cells (hiPSCs) from DMD patients during muscle differentiation offers insights into the regulation of calcium homeostasis. The primary aim of this study was to characterize the pathology in muscle cells derived from DMD patient hiPSCs (hiPSC-skMCs). This characterization involved identifying physiological changes and evaluating the cells’ ability to regulate calcium concentrations at rest and upon stimulation. We subsequently conducted a comprehensive series of tests to assess resting calcium signatures, calcium levels, elevations in calcium following electrical or pharmacological stimulation, and calcium influxes. This study primarily focused on two stages of muscle cell differentiation: the myoblast stage (D3) and the myotube stage (D7). Most of our experiments highlighted calcium dysregulation in DMD hiPSC-skMCs compared to healthy cells. However, our DMD hiPSC-skMCs models did not exhibit calcium dysregulation via purinergic receptors or store-operated channels.

We investigated the resting membrane potential of our cells to assess their excitability and, consequently, their contractile capacity. Our findings showed membrane potential values of −9.0 ± 1.0 mV for myoblasts, −24.0 ± 1.9 mV for myotubes, and −32.8 ± 3.0 mV for myotubes after 7 days of differentiation. In vivo studies in rats have indicated that skeletal muscle cells exhibit a more polarized membrane potential with potential values ranging between −80 and −85 mV (De Jonge WW. et al., 1985). Research on humans has reported similar resting membrane potential values, ranging from −92 to −97 mV (Ruff and Whittlesey, 1992). Our results suggest that hiPSC-skMCs are less polarized, consistent with prior observations on cultured human cells (Iannaccone et al., 1987), which displayed membrane potential values of −27 ± 1 mV for mononucleated myoblasts and −33 ± 2 mV for young myotubes (9 days in vitro). Despite the observed low membrane polarization, we assessed excitation-contraction coupling in our cells using both electrical stimulation and acetylcholine perfusion. Only a small proportion of myoblasts responded to either electrical stimulation or acetylcholine perfusion in both healthy and DMD cells. However, this proportion increased in myotubes, indicating a maturation process of the hiPSC-skMCs. Moreover, when nifedipine was added during acetylcholine perfusion, there was a reduction in the response of over 80% of cells (Figure 6D). Electrical stimulation in the presence of nifedipine resulted in a complete absence of calcium increase in both cell lines (Figure S2). These findings imply that the calcium increases observed during both electrical and chemical stimulations are mediated via DHPR channels.

In skeletal muscle cells, the cytosolic calcium concentration is typically in the range of ∼50-250 nM (Gehlert et al., 2015). For some time, it has been recognized that calcium handling is disrupted in dystrophin-deficient cells. Specifically, intracellular calcium concentrations have been observed to be elevated in mdx mouse fibers (Turner et al., 1988), mdx myotubes (Hopf et al., 1996), and in primary cultures derived from DMD patients (Imbert et al., 1995). In our study, we analyzed intracellular calcium levels to compare healthy and DMD cells derived from hiPSCs (Figure 2A). The results indicated that the intracellular calcium level in healthy cells, both at the myoblast and myotube stages, had an approximate mean value of 120 nM. In contrast, DMD myotubes did not show significant differences when compared to healthy myotubes. However, DMD myoblasts exhibited significantly elevated levels, reaching 290 nM. One potential explanation for these observations is the restoration of resting calcium homeostasis in DMD cells at the myotube stage. This restoration might be attributed to an enhanced storage capacity in hiPSC-skMCs during the myotube stage, possibly due to the maturation and fusion of myoblasts (Ezerman and Ishikawa, 1967). Alternatively, the elevated calcium levels might be associated with increased expression of calcium extrusion pumps (e.g., SERCAs, PMCAs) and exchangers (e.g., NCXs) at the myotube stage. Supporting this hypothesis, our RNAseq analysis revealed heightened mRNA transcript levels for SERCA2 (DMD myoblasts: 41907 ± 4516; DMD myotubes: 71608 ± 5154) and NCX3 (DMD myoblasts: 335; DMD myotubes: 630) (Mournetas et al., 2021). Analysis of spontaneous calcium activity revealed an increase in the amount of calcium released during calcium oscillation throughout differentiation, consistent with observations in mdx mice (Campbell et al., 2006). Additionally, previous research has indicated compromised intracellular calcium cycling in DMD cells (Balghi et al., 2006; DiFranco et al., 2008). When comparing healthy and DMD cells, we observed dysregulation in the amount of calcium released during global spontaneous calcium releases (Figure 3C), aligning with studies on the mdx model (Burr and Molkentin, 2015). Examination of local increase events in hiPSC-skMCs revealed a significant increase in the density of spontaneous calcium release in DMD cells (Figure 4B). Moreover, the kinetics of these events differed between the two cell lines, with DMD cells exhibiting prolonged duration and rise time. An increased density of spark-release sites could indicate dysregulation of ryanodine receptors (RYR) (Pouvreau et al., 2007) and/or IP3 receptors (IP3R), as we have previously shown in dystrophin-deficient muscle cell lines (Balghi et al., 2006; Mondin et al., 2009). The observed kinetic differences might also be attributed to diminished SERCA expression. Dysregulation of SERCA activity has been documented in the mdx mouse model, underscoring the role of SERCA2a in calcium dysregulation in DMD (Goonasekera et al., 2011). Such dysregulation was also confirmed in human cells (Wasala et al., 2020). In line with these findings, reduced mRNA transcript levels were found for SERCA2 in DMD cells compared to healthy cells, as determined by RNAseq analysis, in both myoblasts (Healthy myoblasts: 70138 ± 7142; DMD myoblasts: 41907 ± 4516; p<0.01) and myotubes (Healthy myotubes: 109183 ± 9747; DMD myotubes: 71608 ± 5154; p<0.01) (Mournetas et al., 2021).

Previous studies have shown that dystrophin deficiency impacts excitation-contraction coupling (ECC) during skeletal muscle stimulation (Balghi et al., 2006; Friedrich et al., 2004). Our investigations using electrical stimulation revealed heightened calcium amplitude in DMD cells, suggesting potential ECC dysregulation in dystrophic cells (Figure 5 C). This finding aligns with the observed increase in mRNA copy numbers for the cardiac isoform of DHPR (CACNA1C or Cav1.2) in DMD myoblasts and myotubes (Mournetas et al., 2021). Overexpression of the cardiac isoform of DHPR may contribute to ECC dysregulation. Stimulation via acetylcholine depolarization also led to pronounced calcium amplitude in DMD cells. Notably, kinetics of calcium increases remained consistent across both cell lines. Numerous studies have corroborated our observations, showing elevated calcium responses in DMD cells compared to healthy counterparts (Cognard et al., 1993; Marchand et al., 2004) and others). This supports the notion of disrupted calcium excitation-release coupling in DMD. Conversely, some research indicates that electrically-induced calcium release in mature dystrophic muscle fibers is lower than in healthy fibers (Hernández-Ochoa et al., 2015; Woods et al., 2004, 2005), suggesting that calcium dysregulation may diminish as cells mature into muscle fibers. One theory posits that cells with severe dysregulation might not progress to maturity (De Backer et al., 2002). Collectively, these findings underscore the suitability of hiPSC-skMCs as a model for exploring calcium responses to muscle cell stimulation.

Calcium dysregulation has been demonstrated to be associated with heightened membrane calcium entry due to the dysregulation of TRP channels involved in CCEs, SOCEs, and SACs (Gailly, 2012; Sabourin et al., 2009; Vandebrouck et al., 2006, 2007, 2002). SOCE represents a universal mechanism allowing Ca^2+^ to enter cells in response to a decrease in SR Ca^2+^ content (Putney, 1986). In our study, both CCEs and SOCEs were analyzed in our cells to verify the dysregulation of these pathways in DMD hiPSC-derived muscle cells. Specifically focusing on CCEs, calcium entry was observed in both healthy and DMD lines. However, the amount of calcium entering the cytoplasm was greater in DMD cells compared to healthy cells (Figure 7). This observation aligns with our RNA analysis which highlighted an overexpression of channels involved in CCEs (specifically TRPC4 and TRPC6, data not shown) in DMD hiPSC-skMCs (Mournetas et al., 2021).F. Yet, when analyzing store-operated calcium entry, the quantity of calcium entry through this mechanism was found to be negligible and there were no significant differences between healthy and DMD hiPSC-skMCs. This is intriguing given the pronounced differences observed in the DMD animal model (Onopiuk et al., 2015). Several hypotheses may explain these findings. SOCE involves various proteins such as STIM1/2, ORAI1, and TRPCs (Darbellay et al., 2009, 2010). A closer look at mRNA expression revealed that the transcription of ORAI1, TRPC1, and STIM2 was not significantly altered in DMD cells compared to healthy cells (data not shown). However, a study on a primary culture human model indicated that, at the myotube stage, the SOCE slope was more pronounced in both healthy and DMD cells (Harisseh et al., 2013). This discrepancy might relate to the model itself, potentially due to the incomplete cell maturation of hiPSC-skMCs, which might explain the absence of differences observed in the SOCE analysis for DMD cells.

In this study, we also investigated the activation of purinergic receptors, which are triggered by adenosine 5’-triphosphate (ATP) to facilitate the rapid flux of cations including Na^+^, K^+^, and Ca^2+^ (North, 2002). Prior research on purinergic receptors has highlighted the role of P2X receptors in DMD. Specifically, muscle damage might lead to the release of extracellular ATP, which can then activate specific purinergic receptors such as P2X7 on immune cells, contributing to chronic inflammation (Górecki, 2019). Moreover, other studies showed that high concentrations of extracellular ATP induce abnormal Ca^2+^ influx in DMD cells (Róg et al., 2023; Young and Górecki, 2018). However, in our experiments, when DMD hiPSC-skMCs were perfused with BzATP, an ATP agonist, we observed no calcium release in DMD cells. This suggests that this particular pathway might not be active in our cell model.

In summary, our data strongly suggest that DMD hiPSC-skMCs exhibit intracellular calcium dysregulations, consistent with observations in various other models. Based on RNAseq analyses of these cells, our findings spotlight several mechanisms known to be disrupted in other dystrophin-deficient models. However, certain pathways, such as purinergic and store-operated calcium entry mechanisms, were not found to be dysregulated in our DMD hiPSC-skMCs model. This could suggest that the dysregulations observed in animal models might not be as prominent in human models. Given the diverse origins of these dysregulations, we aligned our findings with (i) previous studies that documented calcium dysregulation pathways in both human and animal models, and (ii) RNA sequencing data identifying potential molecular actors of dysregulation with significantly altered mRNA levels in DMD compared to healthy cells (Mournetas et al., 2021). We subsequently developed a scoring system: the number of studies (with human model-based studies given double weight) multiplied by the count of dysregulated transcripts (DMD vs. healthy). Figure S6 presents the scoring results in a pie chart, with segments—each representing a pathway or dysregulation site—color-coded based on alignment with our findings (green: match, orange: no match, grey: undetermined). This visualization underscores pathways corroborated by our hiPSC-skMCs model data, emphasizing their role in DMD calcium dysregulation, while also spotlighting areas either not universally agreed upon or warranting further investigation. Such insights could guide the identification of promising therapeutic targets for DMD treatment. Yet, the lack of certain responses might be attributed to the inherent limitations of our in vitro model. In a more intricate and differentiated system, dysregulations considered non-significant in our study might become evident. Nonetheless, we believe that DMD hiPSC-skMCs serve as a valuable model for studying calcium dysregulation in Duchenne Muscular Dystrophy.

## Experimental procedures

### Cell cultures

#### Human induced pluripotent stem cell-derived skeletal cells

Three healthy (M180, M398, M194) and two DMD (M202 and M418) cell lines have been used (Mournetas et al., 2021). hiPSC-derived skeletal myoblasts were seeded and differentiated towards myotubes using commercial media (Myoblast Cell Culture Medium SKM02, Myotube Cell Culture Medium SKM03, AMSbio). Myoblasts were thawed and seeded at 8 000 cells/cm² in SKM02 for amplification, then seeded on glass support (glass-bottom 35mm Petri dish, 12 mm, and 30 mm coverslips) pre-coated with carbon and coated with Matrigel (Corning) at a density of 30 000 cells/cm² in SKM02. After 3 days of proliferation (corresponding to myoblasts: D3), cells were cultured with the differentiation medium SKM03 for 4 days (myotube stage: D7). Experiments were performed at D3 and D7 stages.

#### Sol cell line

SolC1(−) were derived from the Sol8 cell line obtained from primary culture of normal C3H mouse soleus muscle (Marchand et al., 2004). Cells were thawed and seeded to obtain 80% confluence in HamF12/DMEM (1:1) medium supplemented with 10% FCS and 1% antibiotics. To induce differentiation, the growth medium was changed to a fusion medium (DMEM supplemented with 2% heat inactivated horse serum, insulin [10 μg/ml, Sigma-Aldrich] and 1% antibiotics). Experiments were performed at the myotube stage, 4 days after addition of fusion medium.

### Intracellular calcium concentration

Intracellular calcium concentration ([Ca^2+^]_i_) was measured using the Fura-2-AM calcium probe (ThermoFisher Scientific). Healthy and DMD hiPSC-derived muscle cells at D3 and D7 were loaded with Fura-2 AM (3 μM) for 30-45 min at 37 °C with 5% CO2. Cells were washed twice with Tyrode’s solution containing 130 mM NaCl, 5.4 mM KCl, 0.8 mM MgCl_2_, 1.8 mM CaCl_2_, 10 mM HEPES, 5.6 mM glucose (pH 7.4). Petri dishes were mounted in the observation chamber and cells bathed in Tyrode’s solution were imaged as previously described (Chapotte-Baldacci et al., 2020). Ratiometric calcium imaging was performed with an Olympus IX73 inverted microscope (Olympus, Tokyo, France). Cells were excited at 340 and 380 nm using a Lambda 421 beam combiner (Sutter Instrument, Ballancourt-Sur-Essonne, France) and the emitted signal acquired at 510 nm using an Andor Zyla 4.2 PLUS cooled sCMOD camera (Andor Technology, Oxford Instruments, Belfast, UK). Images were acquired at 340 nm and 380 nm at a rate of 1 image/s under successive perfused solutions to obtain the minimal and maximal Fura-2 fluorescence ratios. Cells were perfused with Tyrode solution warmed at 37°C to obtain basal calcium fluorescence. Then, Tyrode solution containing 1 mM EGTA (Sigma-Aldrich), 20 µM ionomycine (Sigma-Aldrich) and no calcium was perfused to obtain the minimum [Ca^2+^]_i_ and cells were then perfused with tyrode solution containing 5 mM Ca^2+^ to obtain the maximal [Ca^2+^]_i_. To determine basal [Ca^2+^]_i_, we used Grynkiewicz equation : [Ca^2+^]_i_=Kd*β*(R-Rmin/Rmax-R) as described previously (Grynkiewicz et al., 1985).

### Membrane potential measurements

Electrophysiological studies were realized at day 3, 7 and day 10 on healthy and DMD lines. Tested cells are selected by eye based on their size, their length and their cylindrical with fusiform aspect. Microelectrode measurements were carried out on monolayers of healthy and DMD hiPSC-derived myoblasts and myotubes and the experiments were realized at room temperature (≈ 21°C). Microelectrodes (50-80 MΩ) were pulled from borosilicate glass capillaries with filaments (BF150-110-10, Sutter Instruments) using a horizontal micropipette puller (P-97, Sutter Instruments). The microelectrodes were filled with 3 M KCl and the bath solution was composed of (mM): 150 NaCl, 4 KCl, 1.1 MgCl2, 2.2 CaCl2, 10 HEPES, 5.6 glucose at pH 7.4 adjusted with NaOH solution. Microelectrode is connected to an impedance adapter (H180, Biologic, Claix, France) linked to an amplifier (VF-180, Biologic, Claix, France) and resting membrane potential (RMP) was digitalized with an analog-digital converter (Axon Digidata 1550a, Molecular Devices, San Jose, CA, USA) using pClamp software v10.2 (Molecular Devices, San Jose, CA, USA).

### Spontaneous calcium fluorescence recording

For the characterization of spontaneous calcium activity of healthy and DMD hiPSC-derived myoblasts and myotubes, cells were incubated with Fluo-8 AM calcium dye (5 µM, Abcam) for 15 min at room temperature in Tyrode solution (130 mM NaCl, 5.4 mM KCl, 0.8 mM MgCl_2_, 1.8 mM CaCl_2_, 10 mM HEPES, 5.6 mM glucose, pH 7.4). Fluo-8 fluorescence was recorded with a revolution imaging system from Andor technology feature a spinning disk unit (CSUX1, Yokogawa, Tokyo, Japan) and an Andor Ixon+897 back illuminated EMCCD camera (16µm2 pixel size) on an Olympus inverted IX81S1F-ZDC microscope (Olympus, Tokyo, Japan). IQ3 acquisition software (Andor technology, UK) is used to acquire images of 512×512 pixels. Fluorescence images were visualized through an objective x20 (NA 0.75). Fluo-8 was illuminated at 488 nm with an argon laser and fluorescence emission was collected with a GFP filter set (dichroic mirror: 488 nm, emission filter: 525 nm). Recordings were performed with a x40 objective at an acquisition speed of 5 frames/s for 3 min.

From fluorescence image series, fluorescence measurements were extracted under ImageJ software (NIH, Bethesda, MD, USA). Fluorescence changes were expressed as the fluorescence normalized to basal values (ΔF/F0). Calcium response parameters as amplitude peak (ΔF/F0), area (U.A), duration (s), time to peak (TTP, s), recovery time (RT, s), event frequency (events/min) and activity time were analyzed with a macro-software developed under the image-processing IDL 6.1 language (Research Systems Inc., Boulder, Colorado).

### Localized calcium release analysis

#### Density of release sites

Sequences of images in fast mode were analyzed to calculate the standard deviation of the recorded fluorescence in each pixel as a function of time. During the acquisition sequence, when several releases were observed in the same location, the calculated standard deviation of pixels in this location was higher than in areas without calcium increase. Standard deviation was performed by ImageJ software to obtain one image displaying all fluorescence changes allowing to detect spontaneous localized calcium release sites (Balghi et al. 2006b). Calcium release site density (10 events/µm²) was obtained by counting the number of calcium release site related to cell surface area.

#### Ca^2+^ release events

hiPSCs-derived myoblasts and myotubes were incubated with Fluo-8 and Ca^2+^ release events were recorded under confocal line-scanning microscope at room temperature in Tyrode solution. Fluorescent images were recorded by using a FV-1000 system mounted on an Olympus IX81 inverted microscope (Olympus, Tokyo, Japan). Cells were excited with a 488 nm laser at x40 objective and two-dimensional fluorescence images were acquired in line scan mode (x, t images: 1 line/1,3 ms, line: 100 pixels) with Fluoview software (Olympus, Tokyo, Japan) and analyzed as described previously (Lorin et al., 2013). These recordings were analyzed using the HARVELE software that first allows an automated detection of signals in x, t images and second calculates morphological parameters of each detected event (Sebille et al., 2005). In brief, four parameters were determined for each Ca^2+^ release event: amplitude (ΔF/F0), RT (ms), full width at half maximum (FWHM, µm), and full duration at half maximum (FDHM, ms).

### Electrical Field Stimulation-evoked calcium activity

Healthy and DMD hiPSC-derived myoblasts and myotubes were loaded with Fluo-8 and the recordings were carried out at room temperature in Tyrode solution with the x20 objective under confocal spinning disk microscope as described previously in Spontaneous calcium fluorescence recording section. Electrical stimulation electrodes were placed 1 mm above the cells and are composed of 2 parallel platinum wires connected to an STM4 electrical stimulator (Bionic Instruments). 30 second recordings are acquired at a rate of 1 frame/100 ms using a 488 nm excitation laser. After 10 seconds of recording, cells were stimulated with a single pulse of 1 ms at 100 mA. The recordings are then saved in .tiff format and analyzed under ImageJ and IDL 6.1 software. For each EFS-evoked calcium release, five parameters were determined: amplitude (ΔF/F0), area (A.U.), duration (s), TTP (s) and RT (s).

### Pharmacological-evoked calcium activity

Healthy and DMD hiPSC-derived myoblasts and myotubes were loaded and analyzed as described previously in EFS-Evoked calcium activity section. 3 minutes were recorded at a rate of 1 frame/ 200 ms using a 488 nm excitation laser. After 20 seconds of recording, 100 µM of acetylcholine, 100 or 300 µM of BzATP were perfused in the dish.

### Constitutive calcium entries (CCEs)

CCE were determined by using Fura-2 quenching method as previously described (Chapotte-Baldacci et al., 2020). In brief, healthy and DMD hiPSC-derived myoblasts and myotubes loaded with Fura-2 were bathed in Tyrode solution for 1 min before perfusing manganese solution containing (in mM): 130 NaCl, 5.4 KCl, 0.8 MgCl2, 0.5 Mn^2+^, 10 HEPES, 5.6 glucose, pH 7.4 for 5 min. Fura-2 loaded cells were excited with a LED at 360 nm and fluorescence emission was collected at 510 nm at a frame rate of 1 image/s with Metafluor software (Molecular Devices). The quench rate of fluorescence intensity (%/min) was then calculated to evaluate CCE.

### Store-operated calcium entries (SOCEs)

To analyze SOCE (Harisseh et al., 2013), Healthy and DMD hiPSC-derived myoblasts and myotubes loaded with Fura-2 were bathed in Ca^2+^ free tyrode solution with 0.2 mM EGTA. Then, thapsigargin was perfused (1 µM) to empty calcium stores from the cells, activating SOC channels. After stabilization, calcium is reintroduced in the medium with a solution of tyrode (1.8 mM of Ca^2+^). As a positive control, ionomycin (5 µM) has been perfused. Ratiometric calcium imaging was performed as for intracellular calcium determination. The SOCE slope has been calculated to analyze calcium entries due to SOC channels.

### Immunological staining

Cultured cells were stained with an indirect immunofluorescence. Cells were rinsed with DPBS (Dulbecco’s phosphate buffered saline, ThermoFisher Scientific, Massachusetts, USA) and fixed with 4% Formaldehyde for 10 minutes at room temperature. After fixation, cells were permeabilized with PBS-Triton X-100 at 0.2% for 10 minutes at room temperature. After washing cells with PBS, a second step of permeabilization was realized with PBS-Tween 20 at 0.1% for 30 minutes at room temperature. Then, cells were probed overnight at 4°C with anti-α-Actinin (1:400, Sigma-Aldrich, Missouri, USA), and Anti-Myosin heavy chain (1:500, MF20, DSHB, IOWA, USA) in PBS-Tween. Cells were washed and incubated 3 hours with secondary antibodies, Alexa Fluor 555 conjugated goat anti-chicken and Alexa Fluor 647 Donkey anti-mouse (1:500, ThermoFisher Scientific). Cells were mounted in Mowiol medium (Sigma-Aldrich). Samples were examined with a laser scanning confocal microscope (Olympus FV3000, Japan). 405 nm, 561 and 640 nm laser lines were used for excitation of DAPI (4rn,6-Diamidino-2-phenylindole, Sigma-Aldrich), Alexa Fluor 555 and Alexa Fluor 647 respectively. Emission fluorescence was recorded through spectral detection channels between 430-470 nm (blue), 570-620 nm (red) and 650-750nm (far red fluorescence emission). 1024×1024 pixels images were acquired with UPLAPO100X40HR objectives lens and X2 numerical zooming (0.06 µm pixel size).

### Electron microscopy imaging

hiPSC-derived cells were seeded on a polycarbonate membrane (Costar 3413). After 3 days of culture (D3) and 7 days of culture (D7), cells were fixed for at least 60 min with freshly prepared 2.5% glutaraldehyde overnight at 4°C. After fixation, cells monolayers were washed three times in PBS and then fixed in 4% OsO4 for 1 hour. Cells were washed again three times in PBS. The cells were then dehydrated with sequential washes in 50%, 70%, 90%, 95%, and 100% acetone and then embedded in 50:50 mixture of acetone and Epon resin overnight. The following day specimens were placed into a fresh change of pure resin for 4 hours, then embedded in another fresh change of 100% resin, and placed in a 60°C oven for 24 hours for polymerization. The blocks were then ready for section. Ultrathin sections (60-nm) with EM UC7 ultramicrotome (Leica) were collected on copper grids and stained with 2% uranyl acetate in 50% ethanol for 3 min, followed by incubation in lead citrate (0.08 M) for 6 minutes. Samples were analysed with JEM 1010 electron microscope (Jeol) equipped with a Quemesa Camera (Olympus).

### Statistical analyses

P-values were calculated either by two tailed t-tests or ANOVA and completed by adequate post-tests, as indicated in the corresponding figure legends. These statistical analyses were performed using the GraphPad Prism 8 software.

## Acknowledgements

We would like to thank the ImageUP platform, especially Anne Cantereau and Emile Béré, for their imaging technical support. Additionally, we want to express our thanks to Jenny Colas and Christophe Magaud for their technical assistances during the project. This study was supported by AFM Telethon (#23205 and #23877).

## Author Contributions

A.D. performed experiments, analyzed the data, and wrote the manuscript. C-A. C-B. performed experiments and analyzed the data and revised the manuscript. L.D. performed electron microscopy experiments. E.M. provided protocols for hiPSC-MCs culture, provided expert support and revised the manuscript. M.B. and M.R. provided expert support to the project. A.C. and C.C. designed the electrophysiological approach. C.P. provided expert support to the project and revised the manuscript. S.S. conceived and coordinated the project, analyzed the data, wrote the manuscript and secured fundings.

## Declaration of interests

The authors declare no competing interests.

## Notes

### Competing Interest Statement

The authors have declared no competing interest.

