## Supplemental Figures for "DMD Human iPSC-derived skeletal muscle cells recapitulate selective calcium dysregulation pathways"

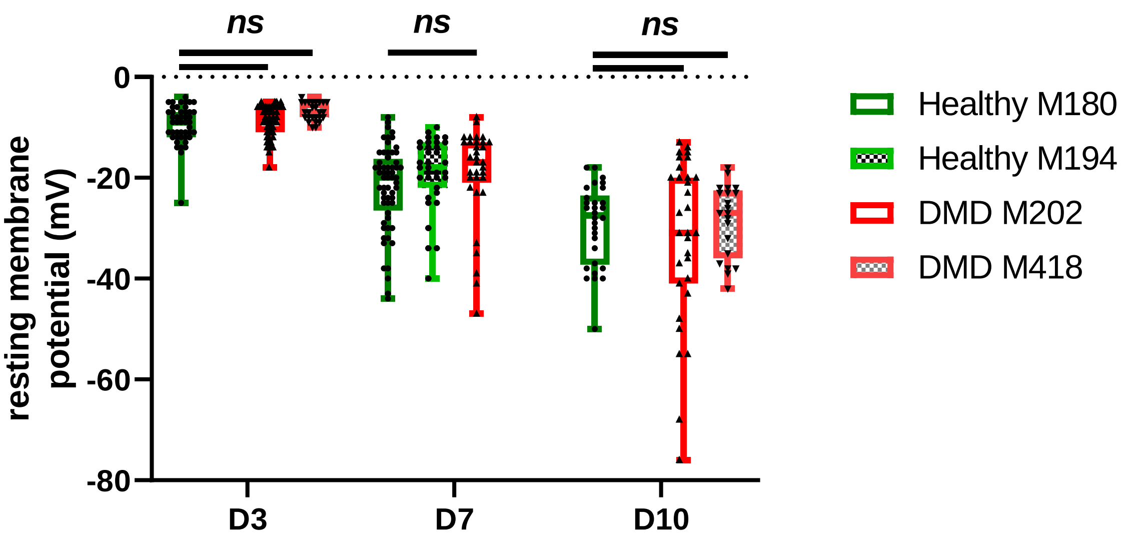


**Figure S1 : Resting membrane potential values in healthy and DMD cell lines.** Resting membrane potential values measured with the microelectrode method in M180 and M194 healthy cell lines (green) and in M202 and M418 DMD cell lines (red) at three different maturation stages, D3, D7 and D10.

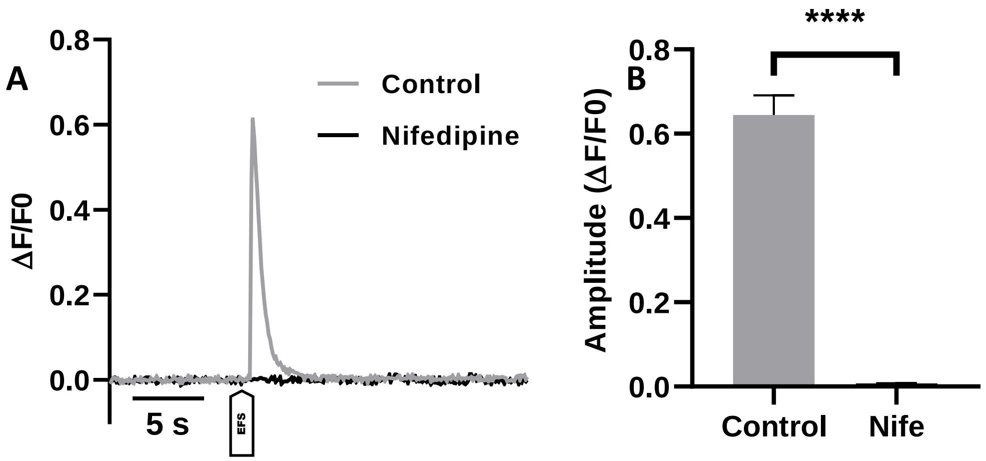


**Figure S2: Characterization of calcium responses during electrical field stimulation with nifedipine incubation.** A: Example of kinetics obtained in DMD (M202) hiPSCs-derived myotubes at D7 with and without nifedipine (1 µM). B: Amplitude (ΔF/F0) of electrically induced calcium increases was determined in DMD hiPSC-CMs at D7 during electrical stimulation. (Data are shown as mean ± SEM. DMD: D7 Control, n=13; D7 Nifedipine, n=87. * indicates the significant difference between DMD cells at D7 without nifedipine to DMD cells at D7 cells without nifedipine. ****; $p < 0.001$, Kruskal Wallis test with Dunn's multiple comparisons test).

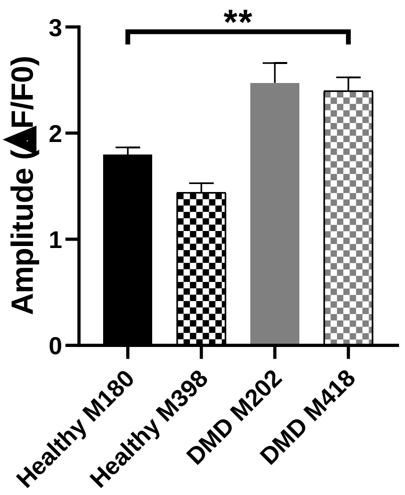


**Figure S3 : Characterization of calcium increase induced by acetylcholine perfusion in DMD and Healthy cell lines at D7.** Analysis of the Amplitude (ΔF/F0) of acetylcholine induced calcium increases in M202 and M418 DMD cell lines and in M180 and M398 healthy cell lines. * indicates the significant difference between DMD and healthy cells at D7. **; $p < 0.01$, Kruskal Wallis test with Dunn's multiple comparisons test).

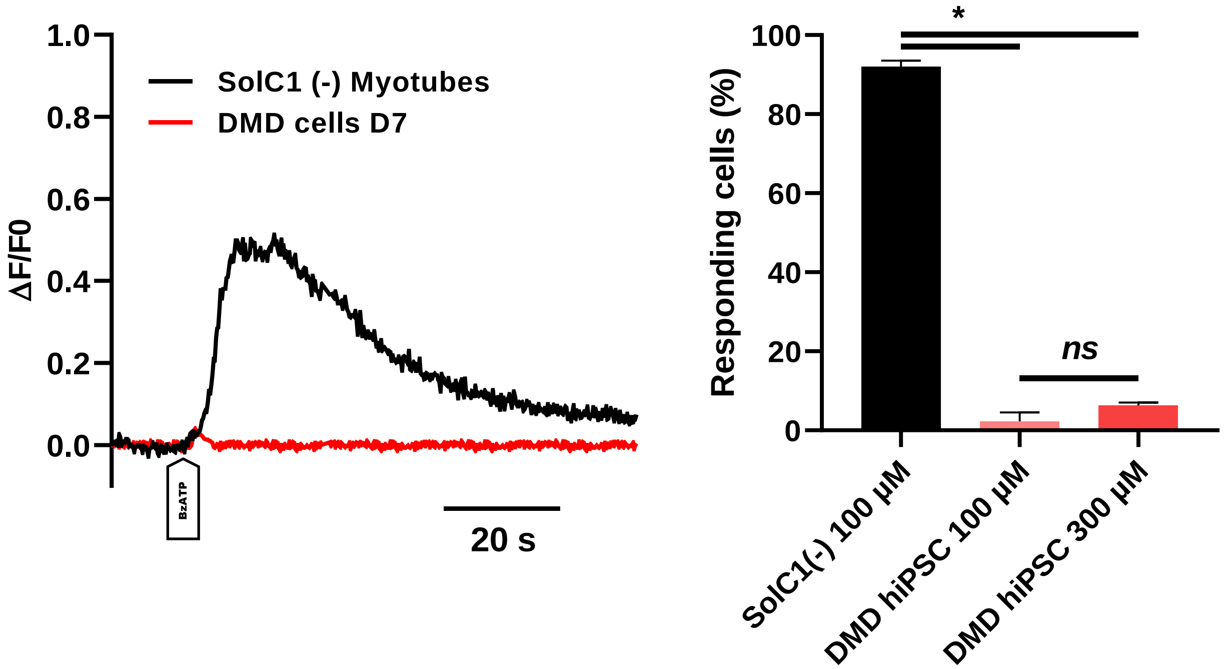


**Figure S4: Characterization of calcium responses induced by BzATP addition in DMD hiPSC-skMCs as compared to dystrophin-deficient mouse muscle cells.** A: Representative traces of calcium kinetics obtained in DMD (M202) hiPSC-skMCs at D7 and in SolC1(-) myotubes. B: Proportion of SolC1(-) and DMD myotubes responding to BzATP stimulation at different concentration (100 µM and 300 µM) SolC1(-) (100 µM), n = 158; DMD hiPSC-skMCs (100 µM), n=40; DMD hiPSC-skMCs (300 µM), n = 43. * indicates the significant difference between SolC1(-) and DMD myotubes. *; $p < 0.05$, Kruskal Wallis test with Dunn's multiple comparisons test).

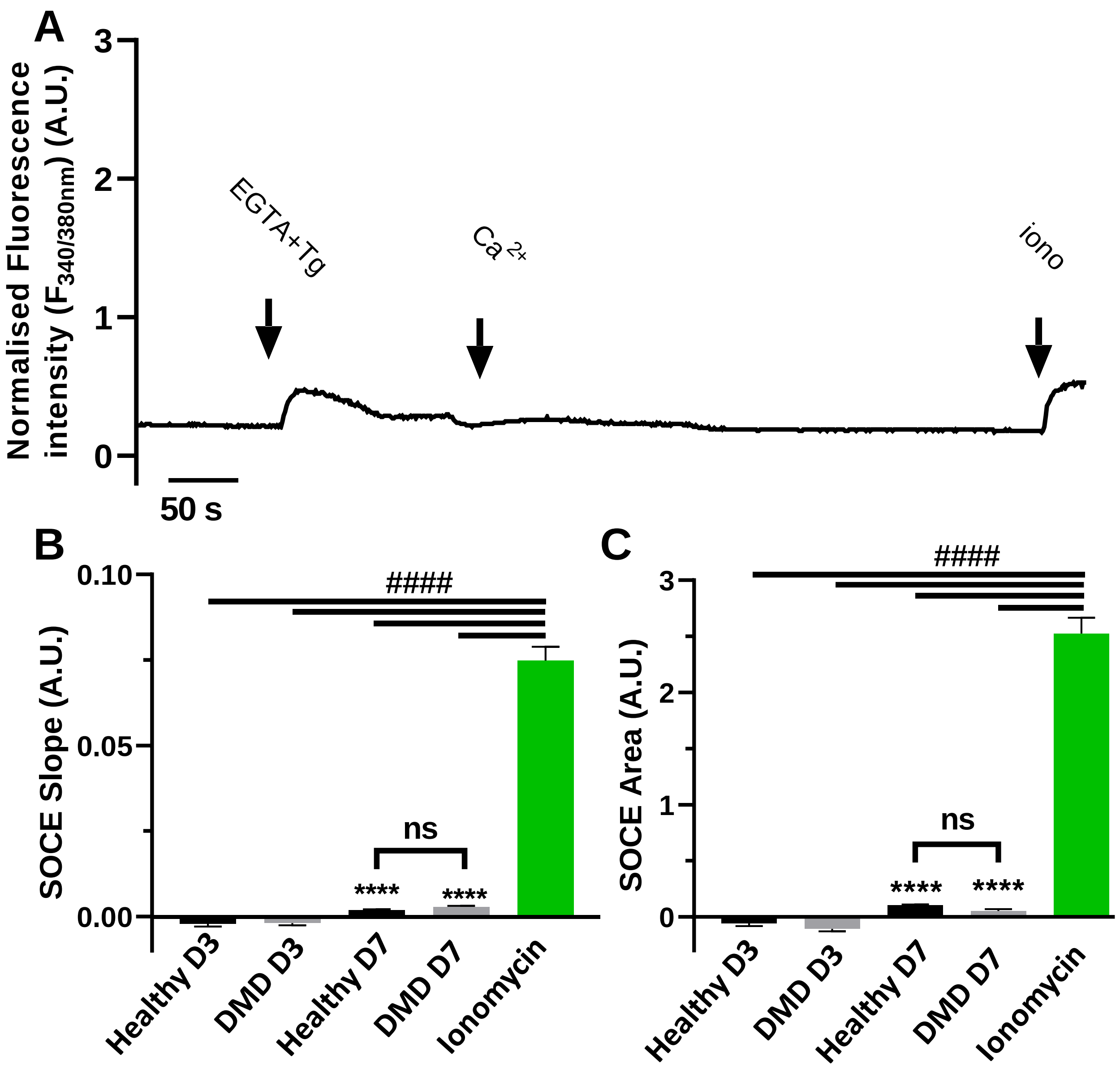


**Figure S5: Characterization of Store-Operated Calcium entries in muscle cells derived from differentiating hiPSCs.** A: Representative traces of calcium kinetics obtained in DMD (M202) hiPSC-skMCs during calcium stores depletion in hiPSCs. B: Measured slopes of calcium increases of both SOCEs when activated with Ca2+ addition and ionomycin when added at the end of the experiment as control. C: measured areas for same signals. # indicates the significant difference between SOCEs and ionomycin (####: $p < 0.001$, Kruskal Wallis test with Dunn's multiple comparisons test.). * indicates the significant difference between D3 and D7 for each line (****: $p < 0.001$, Kruskal Wallis test with Dunn's multiple comparisons test.)

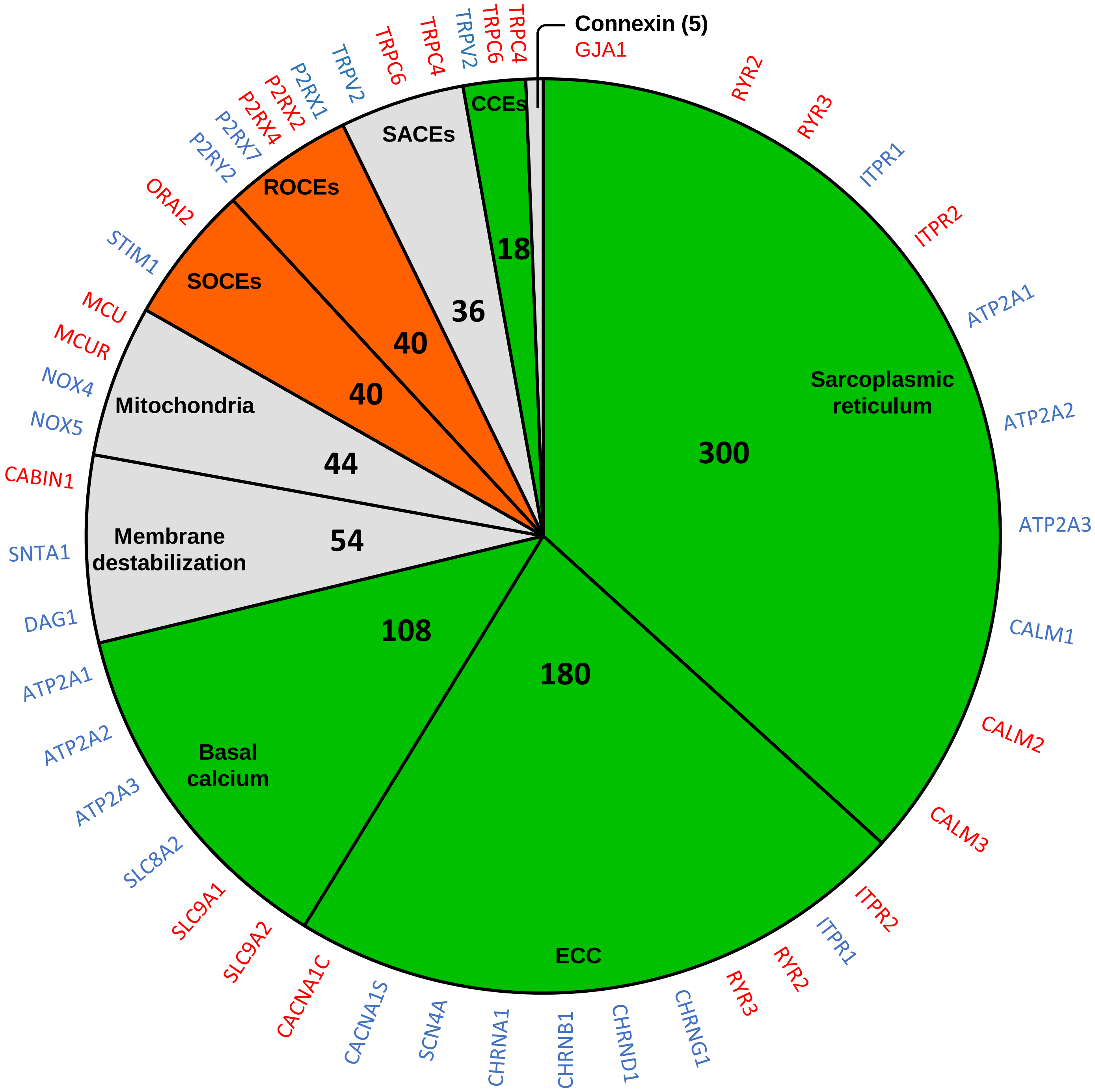


**Figure S6: mRNAs and functional deregulation observed in hiPSCs derived muscle cells at myotube stage.** Pie chart depicting the various origins of calcium dysregulations in DMD. We integrated our findings with (i) previous studies that identified calcium dysregulation pathways in both human and animal models (83 publications, Suppl data S4), and (ii) RNA sequencing results highlighting potential molecular actors of dysregulation. These actors exhibited mRNA levels that were significantly elevated or reduced in DMD compared to healthy controls (Mournetas et al., 2021). We devised a scoring system based on the number of studies, with those conducted on human models given twice the weight, multiplied by the number of dysregulated transcripts (DMD vs. healthy). Each segment, representing a specific pathway or dysregulation site, is both scored and second color-coded based on its alignment with our findings: green for a match, orange for no match, and grey for undetermined. On the pie chart's outer rim, mRNAs of potentially implicated proteins that are notably under-expressed are denoted in blue, while those that are over-expressed are marked in red. Genes without discernible changes in mRNA expression are omitted from the chart. (Mournetas et al., 2021).
